# Potential benefits of Soymilk-*burkina* (*Agbenu*) consumption on gut health of women of reproductive age

**DOI:** 10.64898/2025.12.04.692312

**Authors:** Isaac Darko Otchere, Emmanuel Kyereh, Ethel Juliet Blessie, Isaac Agbemafle, Kwasi Mawuli-Agyenkwa, Kwabena Dentu Gyampo, Akua Boatemaa Arthur, Christopher Galley, Getrude Bubune Ahegbebu, Frank Peget, Nelson Kobla Amey, Richard L.K. Glover, Mary Glover-Amengor

## Abstract

**Background:** Gut microbiome plays a crucial role in human health. This study investigated the impact of two soy-based foods, *soymilk-burkina* (SMB), a traditional fermented Ghanaian beverage) and soymilk-millet blend (SMMB), on the gut microbiome of women of reproductive age.

**Methods:** Fecal samples were collected from two cohorts of women consuming either SMB or SMMB soymilk-*burkina* or a soymilk-millet blend) at 2 weeks pre-intervention, baseline, during an 8-week intervention, and 4 weeks post-intervention. 16S rRNA gene amplicon sequencing was used to analyze the composition and diversity of the gut microbiome.

**Results:** *β*-diversity analysis revealed no significant differences in overall community composition between the two cohorts. *α−*diversity indices (Chao1, Shannon, Simpson) showed no significant differences in richness, evenness, or diversity between cohorts, suggesting a balanced gut ecosystem after both interventions. However, 20 dominant bacterial species were identified, including both beneficial (e.g., *Faecalibacterium prausnitzii* and *Bacteroides vulgatus*) and potentially harmful (e.g., *Eubacterium rectale* and *Bacteroides fragilis*) taxa. *Soymilk*-*burkina* consumption of *soymilk-burkina* significantly reduced the abundance of *Eubacterium rectale* (associated with colon cancer) and increased the abundance of *F. prausnitzii* (beneficial), suggesting potential gut health benefits. However, these effects were transient, highlighting the need for sustained dietary interventions.

**Conclusion:** This study provides insights into the dynamic interplay between diet and the gut microbiome, particularly in the context of traditional fermented foods such as *soymilk-burkina*. Although short-term changes were observed, the long-term benefits may require consistent consumption. Further research is required to explore the mechanisms underlying these effects and their implications on broader health outcomes.

**Plain Language Summary:** The human body, particularly the gastrointestinal tract, hosts trillions of microorganisms, predominantly bacteria. These “gut microbes” play a crucial role in nutrient digestion, vitamin synthesis, and disease prevention. Dietary intake can significantly alter the composition of these microbial communities. This study investigated whether a traditional fermented beverage, *soymilk-burkina*, could enhance the gut health of women in Ghana. A comparative unfermented *soymilk-millet blend* was also examined. Forty women were randomly assigned to two groups, each consuming one of the soy beverages daily for an eight-week period. Fecal samples were collected at baseline, during, and after the intervention to analyze bacterial profiles. The results indicated no dramatic overall changes in the diversity or types of gut bacteria between the two groups. However, women who consumed fermented *soymilk-burkina* exhibited notable shifts, specifically a reduction in *Eubacterium rectale*, a bacterium associated with colon cancer, and an increase in beneficial *Faecalibacterium prausnitzii*, known for its positive impact on gut health. These positive alterations, however, appeared to be transient. Approximately one month after discontinuing the fermented product, the participants’ gut bacterial levels began reverting to their pre-study states. In conclusion, this study suggests that fermented *soymilk-burkina* may offer short-term benefits to gut health by promoting beneficial bacteria and reducing potentially harmful ones. Nevertheless, sustained consumption of this beverage is necessary to maintain these benefits. This research underscores the influence of diet on gut microbiota and highlights the potential of traditional fermented foods like *soymilk-burkina* for further exploration, although more research, particularly concerning long-term effects, is warranted.

## Background

The microbiome of the human body comprises a microbial community, including bacteria, viruses, and fungi, found within and without various organs and tissues. The total number of bacteria alone (38 trillion) is estimated to be greater than the number of our own cells (30 trillion)^1,2.^ Little attention has been paid to these organisms, except for those that pose immediate health problems. However, recently, owing to improvements in detection technologies, attention has been paid to the microbiome of specific organs and tissues of the body, including the gut^3,4^. The human colon (large intestine), which is a part of the gut, is the highest contributor to the microbiome of the body^1^. The gut microbiome is the most diverse and is implicated in a wide range of metabolic processes that are either beneficial (including promotion of fat storage, angiogenesis, training of immunity, biosynthesis of vitamins, metabolism of therapeutics, and breakdown of complex food compounds)^4,5^ or harmful (dysbiosis leading to different disease conditions or directly causing infections) depending on the identity and/or abundance of the organisms^4,5^.

An earlier study reported that simple carbohydrates, including table sugar (sucrose), fruit sugar (fructose), and milk sugar (lactose), do not require digestion after consumption but are readily absorbed in the upper part of the small intestine^6^. However, complex carbohydrates, including starches, cellulose, and fibers, are not easily digested in the stomach and may travel to the large intestine for assisted digestion by enzymes secreted by intestinal microbiota before absorption^6^. Additionally, fermentation of indigestible fibers by intestinal microbes produces short-chain fatty acids that can be used by the body as a source of nutrients and play an important role in muscle function and the prevention of chronic diseases, including certain cancers and bowel disorders^6^. The released short-chain fatty acids lower the pH of the colon, which restricts the range of microbes that can inhabit the colon, limiting the growth of some harmful bacteria, such as *Clostridium difficile*^6^.

The microbiota associated with a healthy gut, including *Prevotella*, *Ruminococcus*, *Bacteroides*, *Firmicutes, Peptostreptococcus, Bifidobacterium, & Lactobacillus* thrive in the hypo-oxygenic colon to prevent the overgrowth of harmful bacteria that enter the body through drinking or eating contaminated water or food by competing for nutrients and attachment sites to the mucus membranes of the gut (a major site of immune activity and antimicrobial protein production)^7–9^. Factors known to directly affect the composition and/or abundance of the gut microbiota include, but are not limited to, the environment, birth mode, use of medications, genetics, and diet, which create a unique person-specific microbiome ^3,10,11^. However, much attention is now focused on the direct impact of diet on shaping the gut microbiome because everyone eats something irrespective of where or how we are born, sick or healthy ^6^. Food substances, including fruits, vegetables, beans, and whole grains such as wheat, oats, millet, and barley, which are rich in prebiotics such as fructo-oligosaccharides, inulin, pectin-resistant starches, and gums, are particularly essential for gut health, as they serve as feed for beneficial microbiota ^12^. However, probiotic foods, including fermented foods, such as *kefir,* yogurt (with live active cultures), pickled vegetables, *tempeh, kombucha* tea, *kimchi, miso*, and sauerkraut, contain beneficial live microbiota that may further alter the microbiome ^13,14^.

Women of reproductive age (WRA) are a special group of people who have a high need for nutrients ^15^ especially during pregnancy and lactation. Women living in poor communities are especially prone to malnutrition, with concomitant health implications ^15^. Therefore, addressing the nutritional needs of this group of women through accessible, nutritious foods holds significant potential for improving maternal and child health outcomes and overall community well-being. However, it is difficult to meet the daily requirements for specific nutrients in communities with poor economic conditions. Women living in poor communities are especially prone to malnutrition, with concomitant health implications ^15^. *Degue,* a fermented cow milk-millet drink containing live organisms from Burkina Faso, has gained popularity in Ghana. It has become a popular source of quick nutrition for both urban and rural dwellers in Ghana. This product is commonly known as *burkina* or *brukina*, because of its origin. The high cost of cow’s milk makes the product quite expensive and unaffordable for the poor, who may depend on it. In addition, this product is not suitable for lactose-intolerant individuals. Our study aimed to use milk produced from soybean (a cheap legume rich in fiber, proteins, magnesium, iron, and fats) as a substitute for cow milk to produce *burkina* and to assess the potential benefits of its consumption on the gut microbiome diversity of WRA in Ghana. We anticipate that this fermented plant-based milk product, called *soymilk-burkina* (SMB), will be cheaper, safer, and more nutritionally beneficial to consumers. Furthermore, in addition to its lower environmental impact compared with rearing livestock for milk, the use of soymilk in *burkina* production will ultimately boost the soybean market, which is being vigorously promoted to enhance the livelihoods of local farmers in Ghana.

## Methods

### Study Area

This study was conducted in Hohoe Municipality in the Volta Region of Ghana (HMV/GH). According to a 2024 report by the Ministry of Finance of the Republic of Ghana (https://mofep.gov.gh/sites/default/files/composite-budget/2024/VR/Hohoe.pdf): HMV/GH has a population size of 114,472 per the 2021 Population Housing Census, with 54,893 being Males (48%) and 59,579 females (52%). The HMV/GH has a high population density (312.3 persons per square kilometer) compared to the regional average (175 persons per square kilometer), with approximately 73% of the population living in urban localities and the remaining 27% living in rural localities. The Municipality has an average household size of 3.1, which is lower than the regional average of 3.3 and a national average of 3.6 individuals.

### Ethics approval

The study protocols and methods were reviewed and approved by the Ghana Health Service Ethics Review Committee (GHS-ERC 015/10/21), Council for Scientific and Industrial Research Institutional Review Board (CSIR/IRB/AL/VOL 1-017) and the University of Health and Allied Sciences Research Ethics Committee (UHAS-REC A7 (2) 2t-22) to ensure integrity of results and protection of participants in accordance with the Declaration of Helsinki.

### Inclusion and Exclusion Criteria

Women of reproductive age (15-49 years), irrespective of pregnancy or breastfeeding, were included in the study, whereas girls aged < 15 years and women aged > 49 years were excluded. Women who were on antibiotics or taking vitamin mineral supplements at least one month prior to the start of the study were also excluded from the study.

### Informed consent and recruitment of participants

Informed consent (read to all WRA, with literates appending their signatures and illiterate thumbprinting) was obtained from all willing WRA before they were recruited for the study. For women below 18 years, their assent in addition to consent by a legal guardian or parent was sought before enrolment. A semi-structured questionnaire was used to capture the demographic information of the recruited WRA. Forty participants, including 10 lactating mothers, 10 pregnant women, and 20 non-pregnant non-lactating (NPNL) women, were recruited for this study (Table 1). The recruited WRA were divided into two feeding cohorts that were fed either the SMB (five lactating mothers, five pregnant women, and 10 neither pregnant nor lactating women) or a soymilk-millet blend (five lactating women, five pregnant women, and 10 NPNL).

**Table 1:**
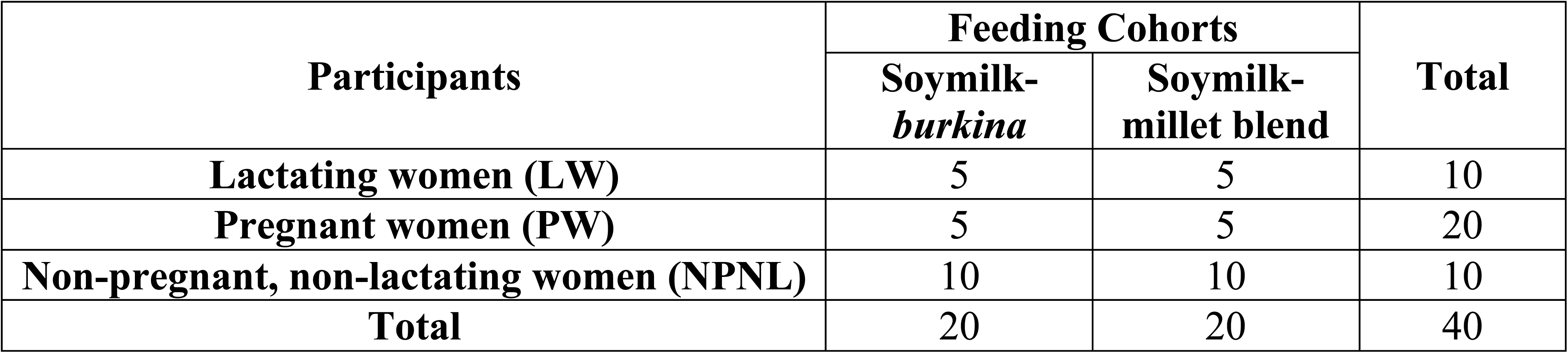
Description and number of study participants.

| Participants | Feeding Cohorts |  | Total |
| --- | --- | --- | --- |
|  | Soymilk-*burkina* | Soymilk-millet blend |  |
| Lactating women (LW) | 5 | 5 | 10 |
| Pregnant women (PW) | 5 | 5 | 20 |
| Non-pregnant, non-lactating women (NPNL) | 10 | 10 | 10 |
| **Total** | 20 | 20 | 40 |

### Production of *soymilk-burkina* (SMB) and soymilk millet blend (SMMB)

The ingredients for the preparation of SMB SMMB included soybeans, pearl millet, sugar, and salt to taste. The production included four distinct stages comprising production of soymilk, fermentation of soymilk into yogurt (for SMB), steaming of ground millet and combination of steamed ground millet with either fresh soymilk or soy yogurt (Supplementary Figure 1).

#### Production of soymilk

Soymilk was produced using an in-house customized procedure. In summary, 2 kg of whole soybeans were steeped in potable water for 8 hours at ambient temperature (28 °C). The beans were subsequently milled into a mash with 6 L of water using an electric grinder. An additional 8 liters (8 L) of water was added before stirring continuously to form a slurry. The slurry was then cooked in a pressure cooker at 110 °C and pressure of 1 bar for 15 minutes. The cooked slurry was filtered to extract 12-15 liters of soymilk. Forty-five grams (45 g) of sugar and 1 g of salt were added per liter of soymilk for improved taste. Vanilla was then added for flavor. A refractometer was used to assess the Brix level, ensuring optimal consistency of the soymilk for each production round.

#### Production of soymilk yogurt

Soy yogurt was prepared following a methodology largely analogous to that of cow milk yogurt^16^. Yogurt starter culture (YC-380, Version: 5 PI EU EN 11-11-2019) was procured from AECI Food & Beverage (a Division of AECI Limited), Johannesburg, South Africa, under license from Chr. Hansen A/S, Denmark. The starter culture was added in proportions of 0.1 g per liter of soymilk and heated to 45 °C with thorough mixing. The inoculated soymilk was then incubated at a pre-set temperature of 45 °C for 8 hours to yield a “drinkable” Soy yogurt.

#### Production of steamed millet agglomerate

One kilogram (1 kg) of millet was washed and steeped for 8 hours. It was then drained, rinsed, and milled into a coarse flour. The coarse floor was sprinkled with water and rubbed between the palms to form raw agglomerates. Fresh water (2L) was boiled over medium heat. The raw agglomerates were to the boiling water and continuously stirred for 20 minutes until a lumpy and sticky consistency was achieved. The lumpy millet (steamed agglomerate) was then removed from the boiling water and allowed to cool at room temperature.

#### Formulation of SMB and SMMB

One liter (1 L) of the soymilk yogurt was measured into a mixing bowl. Six hundred grams (600 g) of steamed millet agglomerate was added and flaked with a fork to break up any lumps then scooped into the mixing bowl containing the soy yogurt. A balloon whisk was used to thoroughly mix the soy yogurt and the steamed millet flakes until the flakes were evenly dispersed throughout the yogurt to form SMB. The formulation of SMMB followed an identical procedure with substitution of unfermented soymilk for soymilk yogurt. The produced SMB and SMMB were refrigerated at 5 °C until ready for consumption.

### Feeding of participants and collection of stool samples

This single-blinded cluster randomized controlled trial was conducted from September to November 2022. The clusters were the different communities that received either SMB or the unfermented product called Soymilk millet blend (SMMB). Each WRA was fed 300 mL of either SMB or SMMB daily by direct observation for eight weeks to ensure compliance. Stool samples were collected using OMNIgene-gut (OM-200) sample collection kits at baseline (pre-intervention week 0), during the intervention period (weeks 6 and 8), and endline (post-intervention weeks 10 and 12), and stored at -80 °C until ready for DNA extraction and metagenomic studies. We chose these pre-intervention, intervention, and post-intervention study designs to ensure that we had clear baseline information before our intervention, detect the potential influence of our intervention, and detect if the influence of our intervention was short-lived or persisted even after the intervention.

### DNA extraction from stool samples for metagenomics analyses

Frozen samples were allowed to thaw at room temperature before DNA extraction under aseptic conditions using a DNeasy PowerSoil ® Pro Kit (Qiagen, Hilden, Germany), according to the manufacturer’s protocol. Briefly, approximately 1000 µL of each liquefied stool sample was added to a Power Bead Pro tube containing lysis buffer and mechanically homogenized for 5 min at maximum speed in a Bead Genie bead beater (Scientific Industries Inc., New York, USA). The inhibitors were removed, followed by the binding of DNA to the solid interphase for washing. Pure DNA was then eluted, and its quality was checked using a NanoDrop 2000 spectrophotometer (Thermo Scientific, Waltham, Massachusetts). The DNA samples were stored at -80 °C until shipment as 60 µL aliquots in sealed 96-well plates at the University of Minnesota Genomics Centre (UMGC), USA, for 16S rRNA V3/V4 region library preparation and sequencing using the Illumina platform according to the manufacturer’s protocols.

### Bioinformatic analysis of sequence

Demultiplexed paired-end sequence reads were trimmed to remove Illumina adapter sequences and filtered to remove low-quality reads using *fastp*^17^. The filtered reads were subjected to taxonomic assignment to various taxa using kraken2 with the standard 32gb kraken microbial database^18^. The relative abundances of the taxonomic assignments were merged into a single. csv files were imported into R for further analysis ^19^. The sample metadata were imported into R and combined with the combined kraken2 report to generate a phylogenetic sequence (phyloseq) object for downstream analyses using the *phyloseq package* of R^20^. Feature tables and data summaries were generated to determine the sequence distribution per sample and *β*-diversity (differences in species composition between two or more samples) using the Weighted UniFrac metric implemented in R, and *α*-diversity (diversity of microbial species within samples) of the samples using the *diversity* function of the *vegan* package in R, after rarefaction with a rarefaction depth of 1000 (below the depth of the sample with the least sequence depth) using the *rarefy_even_depth* function of the *vegan* package of R implemented in RStudio.

### Statistical Analysis

Fisher’s exact test at the 95% confidence level and PERMANOVA analysis were used to carry out *α*-diversity and *β-*diversity analysis between the two groups at specified time points. We used the Bonferroni correction for multiple hypothesis testing to avoid committing a type-1 error (rejecting a true null hypothesis). With an alpha level of 0.05, an adjusted p-value ≤0.01 was considered significant for comparisons between lactating mothers and pregnant women, whereas an adjusted p-value ≤0.005 was considered significant for comparisons between women of reproductive age who were neither pregnant nor lactating.

## Results

### Diversity of gut microbiota

#### Beta (β)-diversity of gut microbiota

We assessed *β*-diversity between the two groups using Weighted Weighted Unifrac metric, which accounts for both phylogenetic relationships and relative abundance of taxa. We found no group-specific clustering of ASV by principal coordinate analysis (PCoA) (Figure 1). The PCoA plots project ASVs onto a two-dimensional space where the distance between points reflects dissimilarity in microbial composition. Samples positioned closer represent similar microbial communities, whereas those farther apart have more dissimilar communities. The PERMANOVA analysis confirmed a statistically insignificant difference in *β*-diversity between groups (p=0.056), indicating no difference in the presence and/or relative abundance of any specific gut microbiota among our study participants, regardless of the intervention and/or time point.

**Figure 1:**
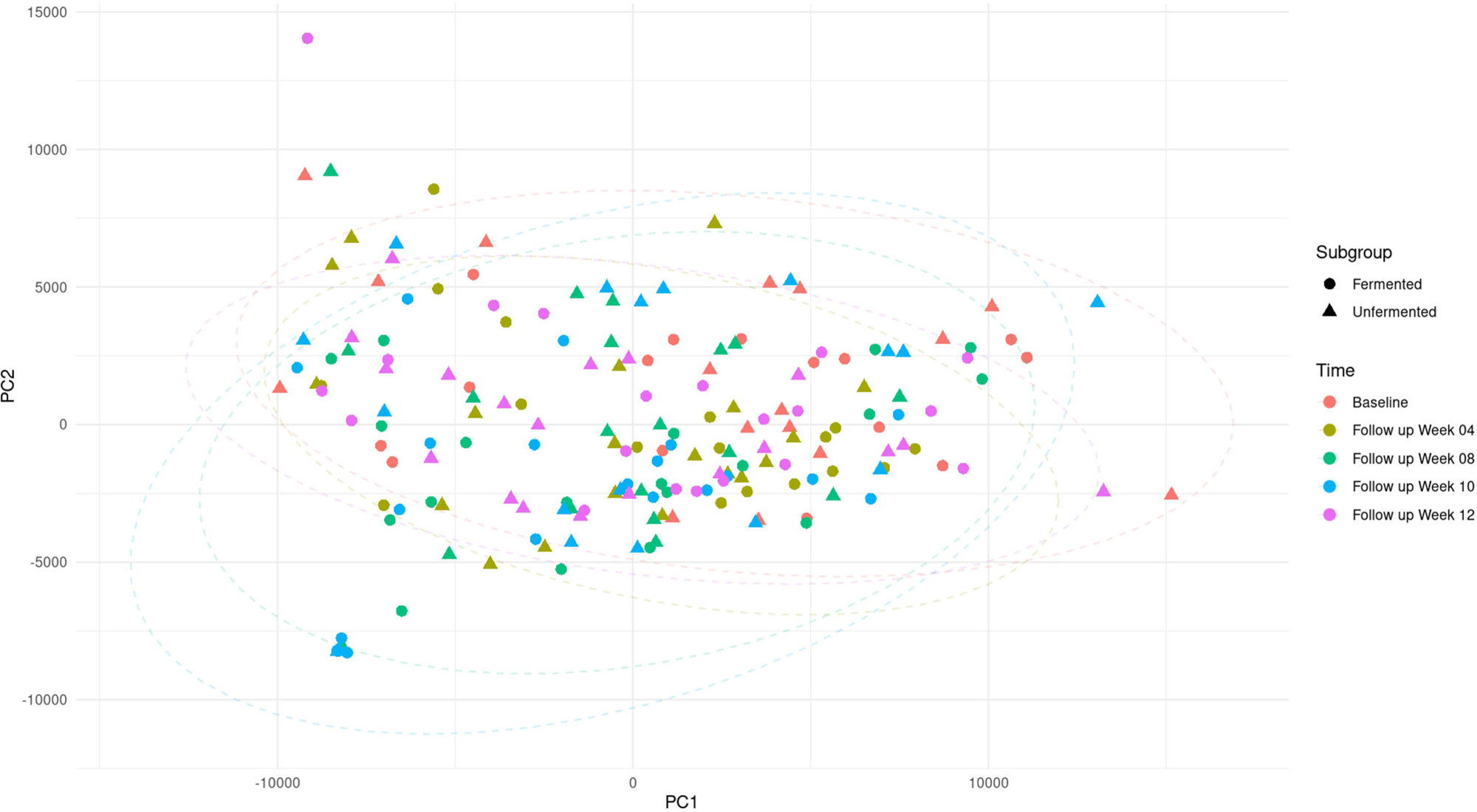
Emperor PCoA plots showing similar distribution of presence and abundance of gut microbiota between the 2 groups. *Dots represent diversity of individual WRAs. Round dots represent WRAs in the SMB arm whereas triangular dots represent those in the SMMB arms. Red, golden-green, green, blue and pink colors respectively represent data points at baseline, week 4, week 8, week 10 (2 weeks post feeding) and 12 (four weeks post feeding)*

#### Alpha (α)-diversity of gut microbiota

We analyzed microbial diversity of the gut microbiome of participants (Figure 2). We found no significant difference in species richness (Chao1 index, p = 0.09) between the gut microbiome of the women in the two groups after four weeks of feeding with SMB and SMMB, albeit the 1^st^ cohort had relatively higher Chao1 index. The difference in species richness between the two groups remained insignificant (p > 0.01) throughout the study period. Similarly, the species evenness (Shannon diversity) and relative abundance of the identified species (Simpson index) were comparable (p > 0.01) between SMB and SMMB after four weeks of feeding. After eight weeks of feeding, there was no difference in the relative abundance and/or evenness of the gut microbiota between the two groups. Although there was a marginal increase in the relative abundance of gut microbiota in the SMB fed cohort, the overall abundance remained insignificant compared to the SMMB cohort (p > 0.01). Furthermore, no significant difference in species evenness and/or relative abundance were observed between the two groups at 4 weeks post-feeding (p > 0.01).

**Figure 2:**
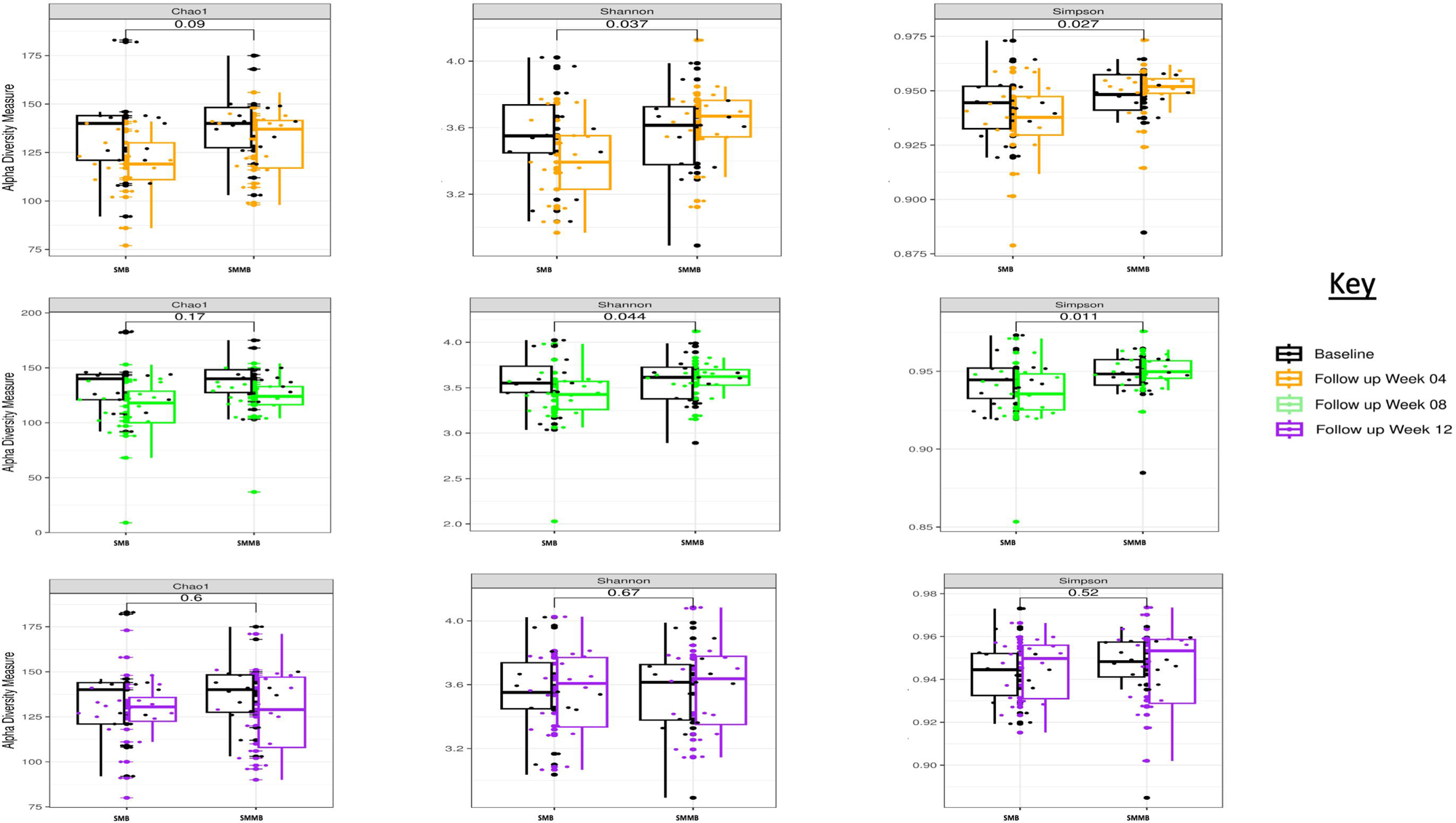
Increasing diversity of gut microbiota among consumers of Soymilk-*burkina and soymilk-millet blend*. *SMB: Soymilk-burkina, SMMB: Soymilk-millet blend*

### Description of the most abundant gut microbiota

In addition to assessing the overall microbial diversity, we analyzed the 20 most abundant bacterial species across all groups and time points. We identified the top 20 bacterial species in the combined dataset and ranked them by their relative abundance as shown below (Table 2). These in order of decreasing abundance were *Faecalibacterium prausnitzii, Prevotella enoeca, Prevotella jejuni, Eubacterium rectale, Eubacterium eligens, Bacteroides vulgatus, Prevotella ruminicola, Oscillibacter valericigenes, Ruminococcus champanellensis, Lachnoclostridium phocaeense, Prevotella dentalis, Collinsella aerofaciens, Blautiasp. N6H1-15, Anaerostipes hadrus, Eubacterium hallii, Dialister sp. Marseille-P5638, Roseburia hominis, Bacteroides helcogenes, Monoglobus pectinilyticus, Ruminococcus bicirculans*. The relative abundance of these species varied between groups before feeding, during the feeding period, and four weeks after feeding, as shown in the heatmap (Figure 3).

**Figure 3:**
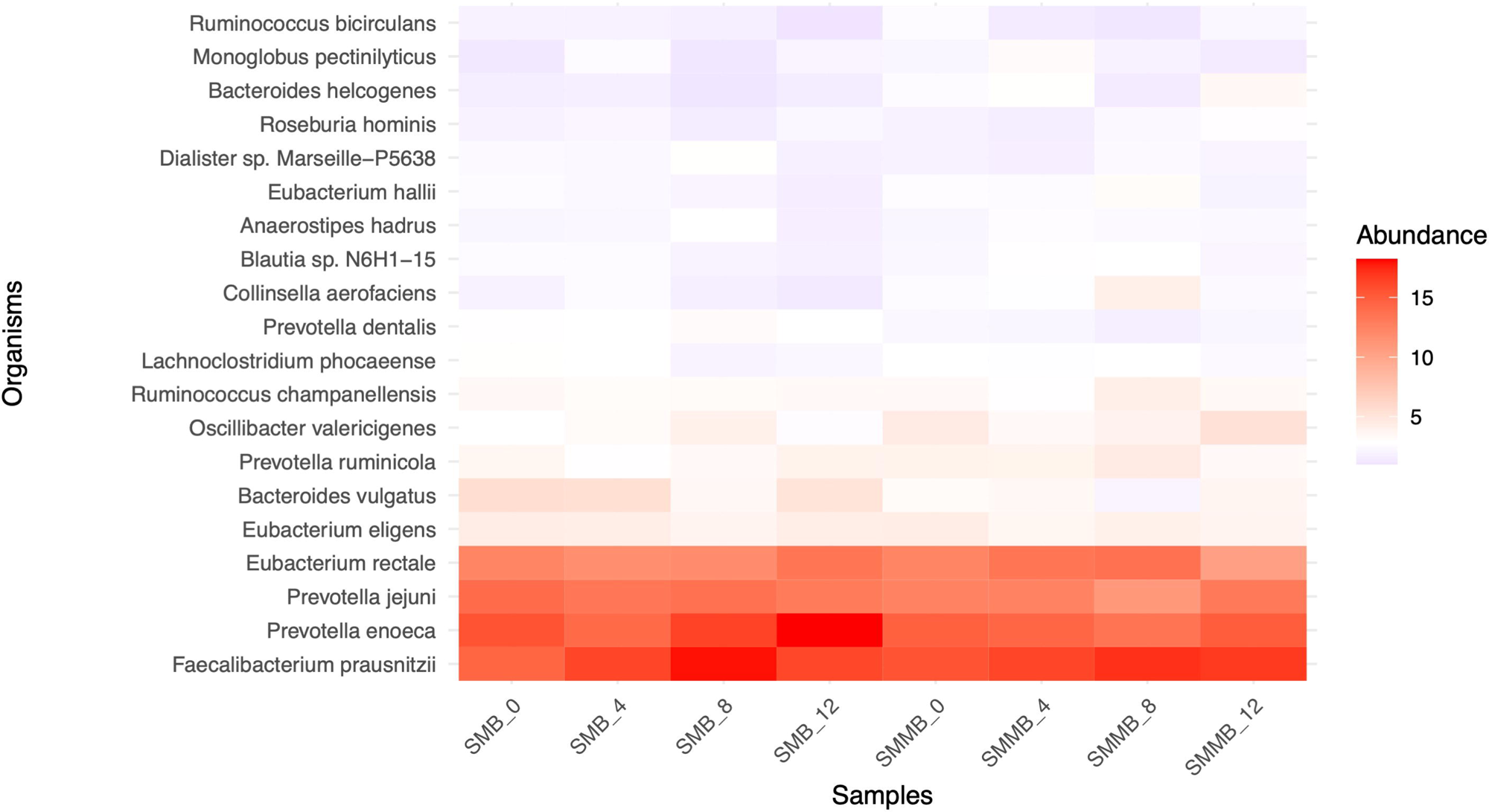
Heatmap of the top 20 most abundant bacteria over the course of the study by cohorts. *SMB = Soymilk-burkina, SMMB = Soymilk-millet blend. WK0 = Before feeding intervention, WK4 = four weeks of feeding, WK8 = eight weeks of feeding, WK12 = four weeks after termination of feeding intervention*.

**Table 2:**
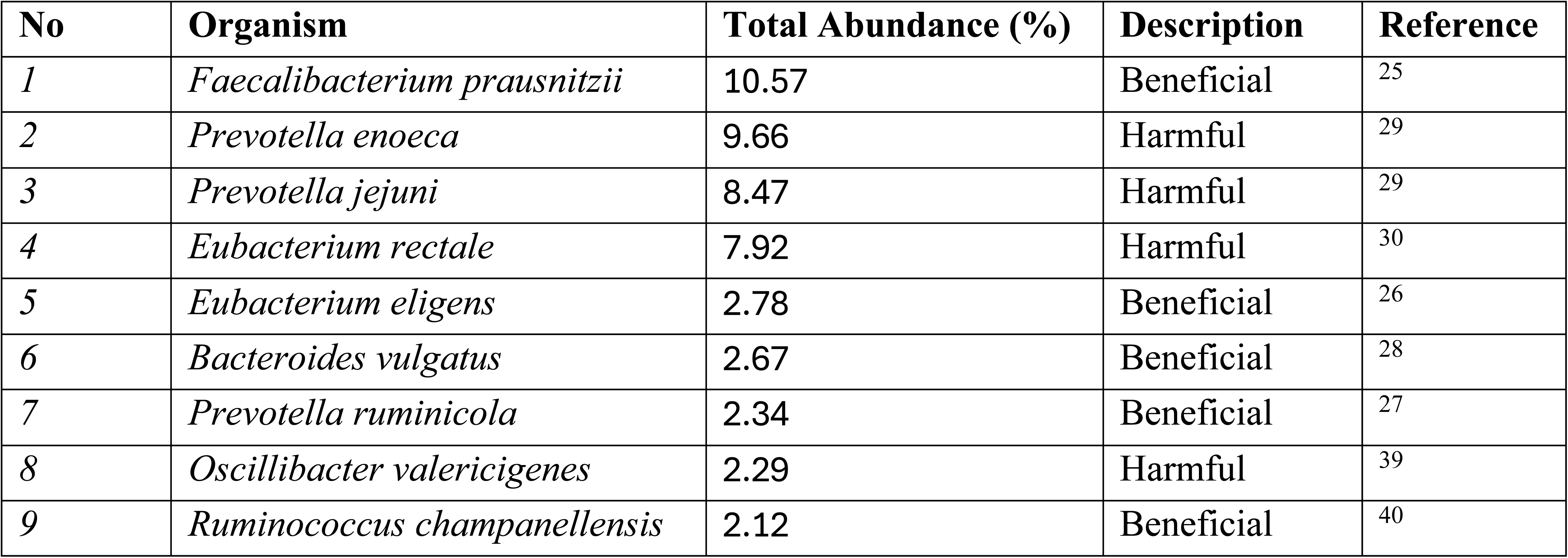
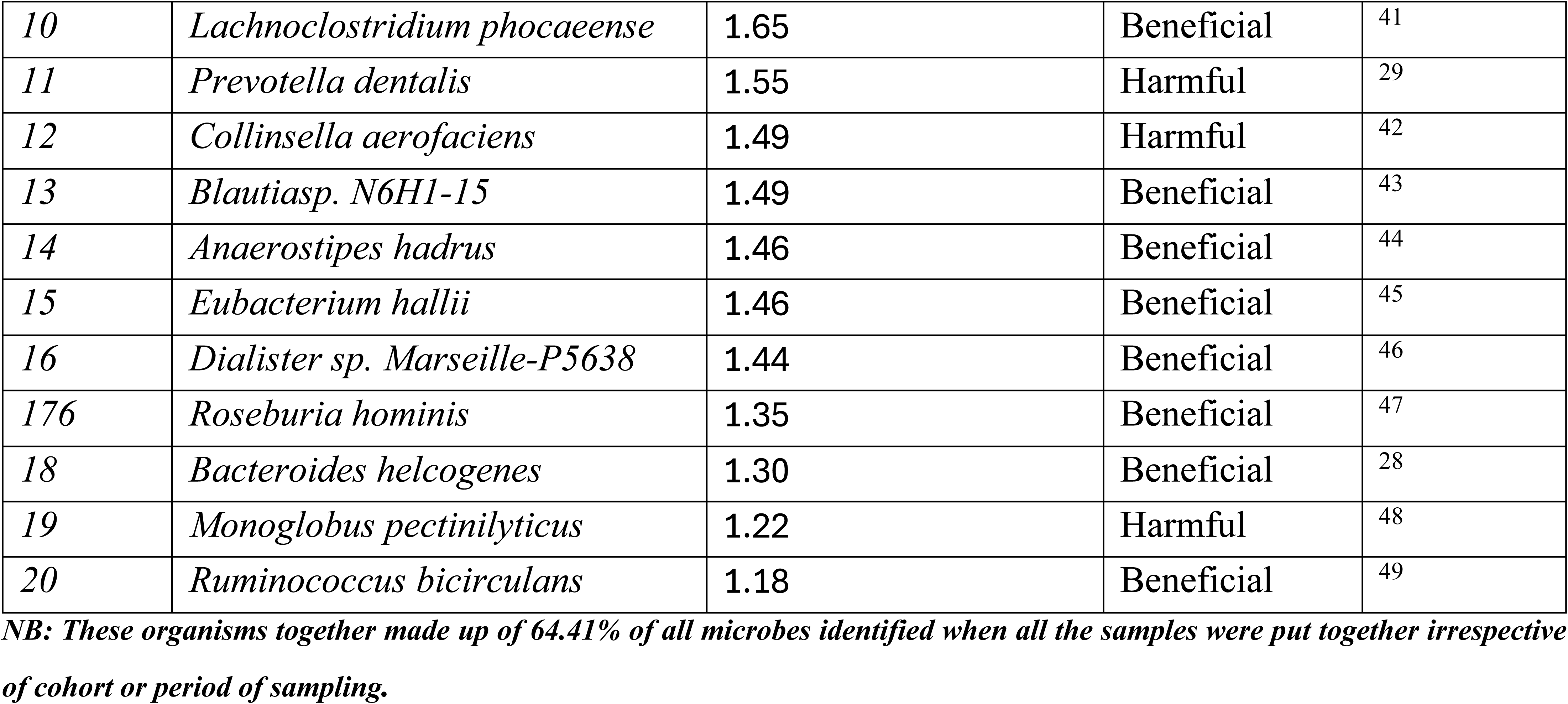
Profile of the 20 most abundant identified bacteria.

| No | Organism | Total Abundance (%) | Description | Reference |
| --- | --- | --- | --- | --- |
| 1 | *Faecalibacterium prausnitzii* | 10.57 | Beneficial | 25 |
| 2 | *Prevotella enoeca* | 9.66 | Harmful | 29 |
| 3 | *Prevotella jejuni* | 8.47 | Harmful | 29 |
| 4 | *Eubacterium rectale* | 7.92 | Harmful | 30 |
| 5 | *Eubacterium eligens* | 2.78 | Beneficial | 26 |
| 6 | *Bacteroides vulgatus* | 2.67 | Beneficial | 28 |
| 7 | *Prevotella ruminicola* | 2.34 | Beneficial | 27 |
| 8 | *Oscillibacter valericigenes* | 2.29 | Harmful | 39 |
| 9 | *Ruminococcus champanellensis* | 2.12 | Beneficial | 40 |
| 10 | *Lachnoclostridium phocaeense* | 1.65 | Beneficial | 41 |
| 11 | *Prevotella dentalis* | 1.55 | Harmful | 29 |
| 12 | *Collinsella aerofaciens* | 1.49 | Harmful | 42 |
| 13 | *Blautiasp. N6H1-15* | 1.49 | Beneficial | 43 |
| 14 | *Anaerostipes hadrus* | 1.46 | Beneficial | 44 |
| 15 | *Eubacterium hallii* | 1.46 | Beneficial | 45 |
| 16 | *Dialister sp. Marseille-P5638* | 1.44 | Beneficial | 46 |
| 176 | *Roseburia hominis* | 1.35 | Beneficial | 47 |
| 18 | *Bacteroides helcogenes* | 1.30 | Beneficial | 28 |
| 19 | *Monoglobus pectinilyticus* | 1.22 | Harmful | 48 |
| 20 | *Ruminococcus bicirculans* | 1.18 | Beneficial | 49 |

NB: These organisms together made up of 64.41% of all microbes identified when all the samples were put together irrespective of cohort or period of sampling.

We searched available literature for the potential impact of the identified twenty most abundant bacteria from our analyses on gut health (Table 2) and identified 13 beneficial bacteria (*Faecalibacterium prausnitzii, Eubacterium eligens, Bacteroides vulgatus, Prevotella ruminicola, Ruminococcus champanellensis, Lachnoclostridium phocaeense, Blautiasp. N6H1-15, Anaerostipes hadrus, Eubacterium hallii, Dialister sp. Marseille-P5638, Roseburia hominis, Bacteroides helcogenes, Ruminococcus bicirculans*) and seven harmful bacteria (*Prevotella enoeca, Prevotella jejuni, Eubacterium rectale, Oscillibacter valericigenes, Prevotella dentalis, Collinsella aerofaciens, Monoglobus pectinilyticus*) according to literature (Table 2).

Using one example each of beneficial (*F*. *prausnitzii* - producer of butyrate in the gut) and harmful bacteria (*E. rectale* - associated with colon cancer) to assess the potential impact of consuming SMB compared to SMMB on gut health, we found that consuming SMB significantly (p<0.0001) reduced the odds ratio (OR) of having *E. rectale* in the gut compared to SMMB from 1.15 to 0.88 (Table 3). In contrast, although SMB had the highest odds of *F. prausnitzii* carriage compared to SMMB (OR=1.07, 95%CI=1.06-1.08, p<0.0001) prior to feeding, there was no significant difference in carriage after the intervention period (OR=0.99, 95%CI=0.99-1.01, p=0.5375).

**Table 3:** Comparing the potential impact of consuming Soymilk-*burkina* on gut health.

| Comparison | Stage | OR | 95%CI | P-value |
| --- | --- | --- | --- | --- |
| SMB: SMMB [*E. rectale*] | Before feeding | 1.15 | 1.14-1.16 | <0.0001 |
| SMB: SMMB [*E. rectale*] | 8 weeks of feeding | 1.15 | 1.14-1.67 | <0.0001 |
| SMB: SMMB [*E. rectale*] | 4 weeks post feeding | 0.88 | 0.87-0.89 | <0.0001 |
| SMB: SMMB [*F. prausnitzii*] | Before feeding | 1.07 | 1.06-1.08 | <0.0001 |
| SMB: SMMB [*F. prausnitzii*] | 8 weeks of feeding | 0.98 | 0.97-0.99 | =0.0002 |
| SMB: SMMB [*F. prausnitzii*] | 4 weeks post feeding | 0.99 | 0.99-1.01 | =0.5375 |

SMB= Soymilk-burkina, SMMB= Soymilk-millet blend.

### Distribution of beneficial and harmful gut microbiota

Prior to the feeding intervention, the total relative abundance of identified beneficial gut microbes was 31.69% and 31.52% for SMB and SMMB, respectively (Figure 4). The abundance of these beneficial microbes increased significantly (p=0.0001) after eight weeks of feeding, peaking at 21.15% and 21.31% for SMB and SMMB, respectively. The proportion of these beneficial microbes reduced significantly (p<0.0001) four-weeks after termination of feeding to 30.14% for SMB but remained higher at 32.40% for SMMB. In contrast, the proportion of known harmful gut microbiota in the SMB was 32.72% before the feeding intervention reduced to 32.31% after four-weeks of feeding, with a further mild reduction to 32.26% at 8-weeks before a sharp increase in abundance to 34.27% four weeks after the intervention (Figure 4). A similar trend was observed for the abundance of harmful gut microbiota in SMMB with 32.89% before intervention, peaking at 33.31% at four-weeks of feeding, significant (p<0.0001) reduction to 32.00% at eight-weeks of feeding, and remaining constant (32.01%) at four-weeks post feeding.

**Figure 4:**
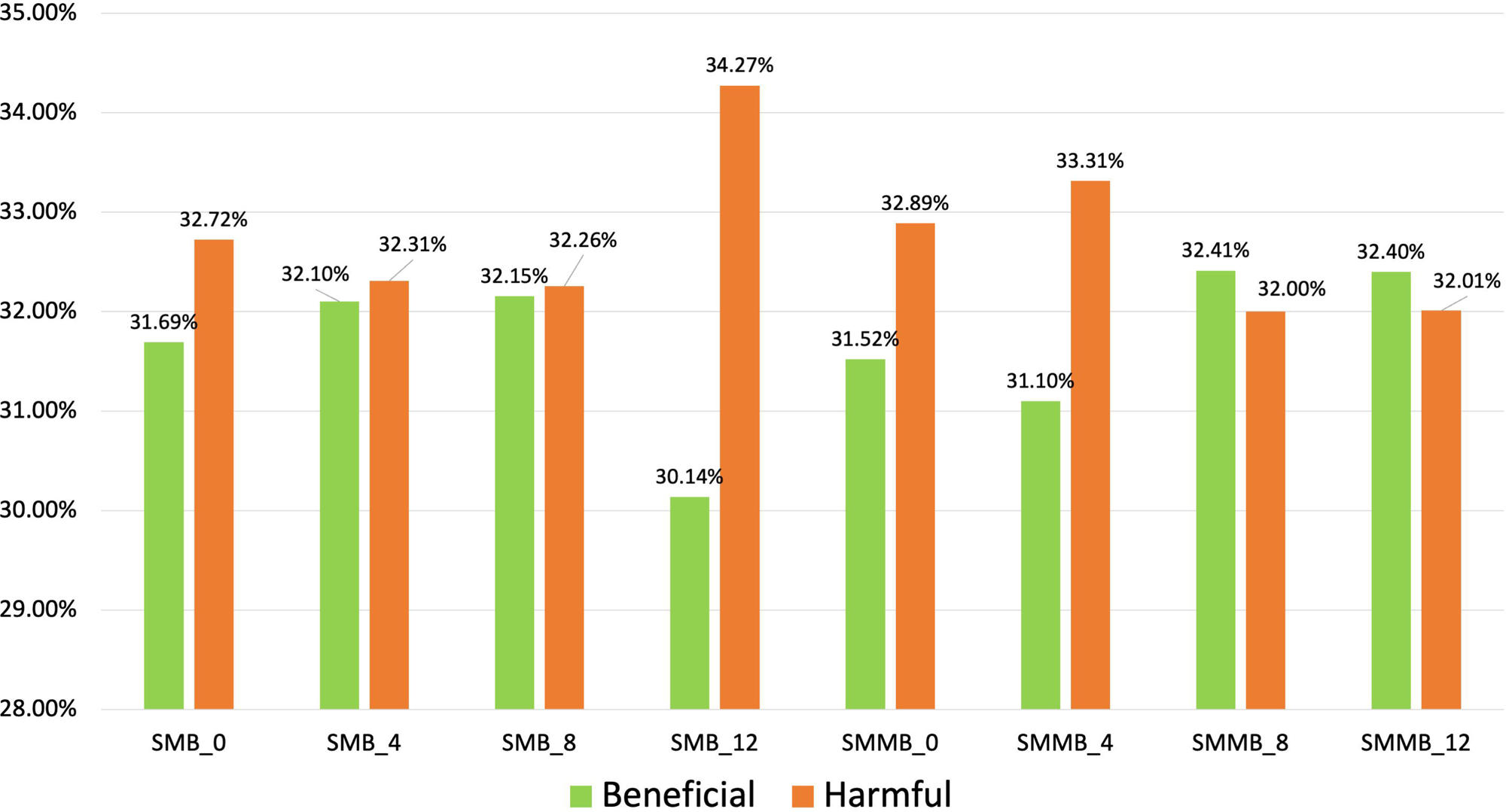
Relative abundance of known beneficial and harmful gut microbes among study cohorts. SMB = Soymilk-burkina; SMMB = Soymilk-millet blend; WK0 = Before feeding; WK4 = four weeks of feeding; WK8 = eight weeks of feeding; WK12 = four weeks after termination of feeding intervention.

## Discussion

The human gut microbiome is a complex ecosystem pivotal to health and disease, and dietary interventions represent a significant avenue for modulating its composition and function^4,5^. Our study specifically aimed to investigate the impact of two soy-based foods, *soymilk-burkina* (SMB), a novel fermented Ghanaian beverage and soymilk-millet blend (SMMB), on the gut microbiome of women of reproductive age (WRA) in Ghana. This objective was driven by the potential for SMB to serve as an affordable, nutritious, and accessible alternative to traditional dairy-based *burkina*, addressing nutritional needs in communities with economic constraints and high nutrient demands among WRA.

We found no significant differences in *β*-diversity (overall community composition) between the SMB and SMMB cohorts. Similarly, *α*-diversity indices (Chao1, Shannon, Simpson) showed no significant differences in species richness, evenness, or diversity. This suggests that, at a broad ecological level, both soy-based interventions promoted a relatively balanced gut ecosystem, a critical aspect for maintaining overall health^21^. This observation, however, stands in contrast to some earlier reports highlighting rapid, broad-spectrum responsiveness of the gut microbiome to diverse dietary inputs^6,12–14^. Some of these earlier reports had demonstrated diet-linked microbiome alterations within short timeframes, contrary to the observations of this study^22–24^. Nevertheless, the contrast of our findings to some of the existing literature suggests that while overall diversity might be maintained, the specific composition at a finer taxonomic level warrants closer examination.

Indeed, our detailed analysis of the 20 most abundant bacterial species (accounting for 64.41% of the sequence reads) unveiled a dynamic interplay between beneficial and potentially harmful taxa, providing a more granular understanding of the interventions’ effects. We specifically noted the presence of health-promoting bacteria such as *F. prausnitzii, E. eligens, P. rumincola* and *B. vulgatus* known for the production of butyrate, anti-inflammatory properties, assisted digestion of plant-rich diets, and contributions to gut barrier integrity respectively ^25–28^. Simultaneously, potentially harmful bacteria as *P. enoeca*, *P. jejuni* and *E. rectale* respectively associated with opportunistic infections, periodontitis, and colorectal cancer^29,30^ were also identified. However, the relative abundance of these microbes flatulated over the study period indicating an active, albeit specific, shifts influenced by the dietary composition^23,31^.

Crucially, when focusing on indicator species relevant to gut health outcomes, our study provided direct evidence supporting the potential benefits of SMB consumption, aligning with our overarching aim. SMB consumption was associated with a significant reduction (p<0.001) in the abundance of *E. rectale*, a bacterium linked to colorectal cancer ^30^, and a concomitant increase (p<0.001) in *F. prausnitzii*, a widely recognized beneficial bacterium vital for gut health ^32,33^. This specific modulation suggests that SMB may actively promote a healthier gut environment by suppressing potentially pathogenic taxa while fostering beneficial ones. This finding is particularly significant given the background rationale of developing a fermented plant-based product with tangible health benefits.

However, a key observation was the transient nature of these beneficial shifts. The positive changes observed during the intervention, particularly the reduction in harmful microbiota and increase in beneficial microbes in the SMB cohort, tended to reverse within four weeks post-intervention. This highlights the impermanence of diet-established gut microbiome changes ^22–24^ and underscores the need for consistent dietary patterns, as previously reported^23,34^. While the short-term benefits are clear, this transient effect emphasizes that continuous consumption of SMB is likely necessary to maintain its protective effects against conditions like colorectal cancer and to provide sustained health advantages. Unfortunately, most of such diets are expensive and unaffordable to vulnerable groups of people ^35–38^. However, SMB, which is prepared from relatively cheap soybeans and millet, can be made relatively cheap for these vulnerable groups to help bridge the nutritional gap.

Our findings contribute to the understanding of how plant-based food interventions, particularly those involving fermentation, can modulate microbial composition. By demonstrating specific benefits from SMB consumption, this study supports the utility of traditional fermented foods for improving gut health, especially within the context of local food systems and affordability. This has significant implications for public health strategies aimed at promoting gut health and overall well-being in WRA and similar populations, especially where access to diverse and expensive probiotic foods is limited. Future research, including larger cohorts and metabolic profiling, is essential to further elucidate the mechanisms and long-term implications of these beneficial dietary effects.

## Limitations and Future Directions

This study is limited by recruitment of participants from a single community in Ghana. This constraint could be mitigated by replicating the study with larger cohorts across multiple communities to confirm its robustness and to investigate whether potential effects vary based on demographics, genetics, or baseline health differences. Secondly, future studies must complement gut microbiota composition analysis with metabolite profiling of stool samples to elucidate the functional consequences of dietary modifications within the gut environment, as changes in microbial abundance do not consistently translate into altered host-microbial metabolic interactions. Furthermore, examining how the observed microbial shifts, particularly the increase in *F. prausnitzii* and reduction in *E. rectale*, correlate with enhanced nutrient absorption from the fiber- and protein-rich SMB and SMMB would be crucial, especially for women of reproductive age who have elevated nutritional demands. Additionally, given the transient nature of the beneficial effects observed in this study, a critical consideration for long-term sustainability, including an understanding of dietary patterns and consistency, is necessary. This will aid in maintaining a resilient gut microbial composition and function, ensuring that affordable solutions like SMB can deliver lasting health benefits in resource-limited settings. Finally, the absence of a control group that consumed no soy-based meals restricted the interpretation of the observed changes. Therefore, future studies should incorporate such control group to assess the specific effects of each intervention compared to a defined placebo group.

## Conclusion

This study aimed to assess the potential benefits of consuming *soymilk-burkina* (SMB), a novel fermented soy-millet beverage, on the gut microbiome diversity of women of reproductive age (WRA) in Ghana, specifically as a more affordable and accessible alternative to traditional dairy-based burkina. Our findings provide valuable insights into the dynamic interplay between this plant-based dietary intervention and the gut microbiota. While overall alpha and beta diversity analyses revealed no significant long-term differences in microbial richness, evenness, or community composition between women consuming SMB and those consuming the unfermented soymilk-millet blend (SMMB), specific and clinically relevant shifts in microbial abundance were observed. Critically, the consumption of SMB significantly reduced the abundance of *Eubacterium rectale*, a bacterium associated with colon cancer, while concurrently increasing the abundance of beneficial *Faecalibacterium prausnitzii*, a key producer of butyrate. These targeted changes suggest a positive modulation of the gut microbiome towards a healthier profile, directly addressing our aim of identifying potential health benefits. However, the transient nature of these beneficial shifts, with microbial compositions reverting towards baseline four weeks post-intervention, underscores the importance of sustained consumption for maintaining a resilient and health-promoting gut microbiome. This research demonstrates the potential of affordable, traditional fermented foods like *soymilk-burkina* to influence gut health, offering a promising avenue for improving nutritional outcomes in vulnerable populations. Further research is warranted to elucidate the precise mechanisms underlying these effects, explore the long-term impact of consistent SMB consumption, and investigate its broader implications for WRA health outcomes in Ghana and similar settings.

## Supporting information

Supplementary Figure 1 ( depicts the SMB and SMMB production pipelines), Supplementary Table 1 (lists the accession numbers of the sequenced samples)

## Data Availability and Sharing

All DNA sequences generated and used in this study with accession numbers (Supplementary Table 1) were deposited in the NCBI database under the accession number SUB13443825. All accompanying metadata can be accessed through a written request to the corresponding author.

## Author Contributions

All authors made a significant contribution to the work reported. whether that is in the conception, study design, execution, acquisition of data, analysis and interpretation; took part in drafting and critically reviewing the article; gave final approval of the version to be published; have agreed on the journal to which the article has been submitted; and agree to be accountable for all aspects of the work.

## Acknowledgement

We appreciate funding from the Bill and Melinda Gates Foundation, without which this study could not have been conducted. Further appreciation goes to the women of reproductive age in the Hohoe Municipality of the Volta Region of Ghana, who volunteered to participate in the study, as well as their families, who supported them throughout the study. We extend our gratitude to the management, administrative, and laboratory staff of the CSIR-Food Research Institute (CSIR-FRI), Accra, Ghana, for their immense contribution, including the formulation of the product, preliminary tests, and administrative support throughout the project. Management of the Fred N. Binka School of Public Health (FNBSPH), University of Health and Allied Sciences (UHAS), Hohoe, Ghana for hosting intervention studies. The Bacteriology Department of the Noguchi Memorial Institute for Medical Research (NMIMR), University of Ghana, Legon, Accra, also deserves special mention for making laboratory facilities and technical staff available for DNA extraction and other confirmatory genomic studies. Our appreciation also goes to the students of the FNBSPH-UHAS, who assisted with the recruitment of participants and administration of food formulations. Funding Statement

This work was supported, in whole or in part, by the Gates Foundation [INV-033569]. The conclusions and opinions expressed in this work are those of the author(s) alone and shall not be attributed to the Foundation. Under the grant conditions of the Foundation, a Creative Commons Attribution 4.0 License has already been assigned to the Author Accepted Manuscript version that might arise from this submission. Please note works submitted as a preprint have not undergone a peer review process

