## Supplementary Figure 1 ( depicts the SMB and SMMB production pipelines), Supplementary Table 1 (lists the accession numbers of the sequenced samples) for "Potential benefits of Soymilk-*burkina* (*Agbenu*) consumption on gut health of women of reproductive age"

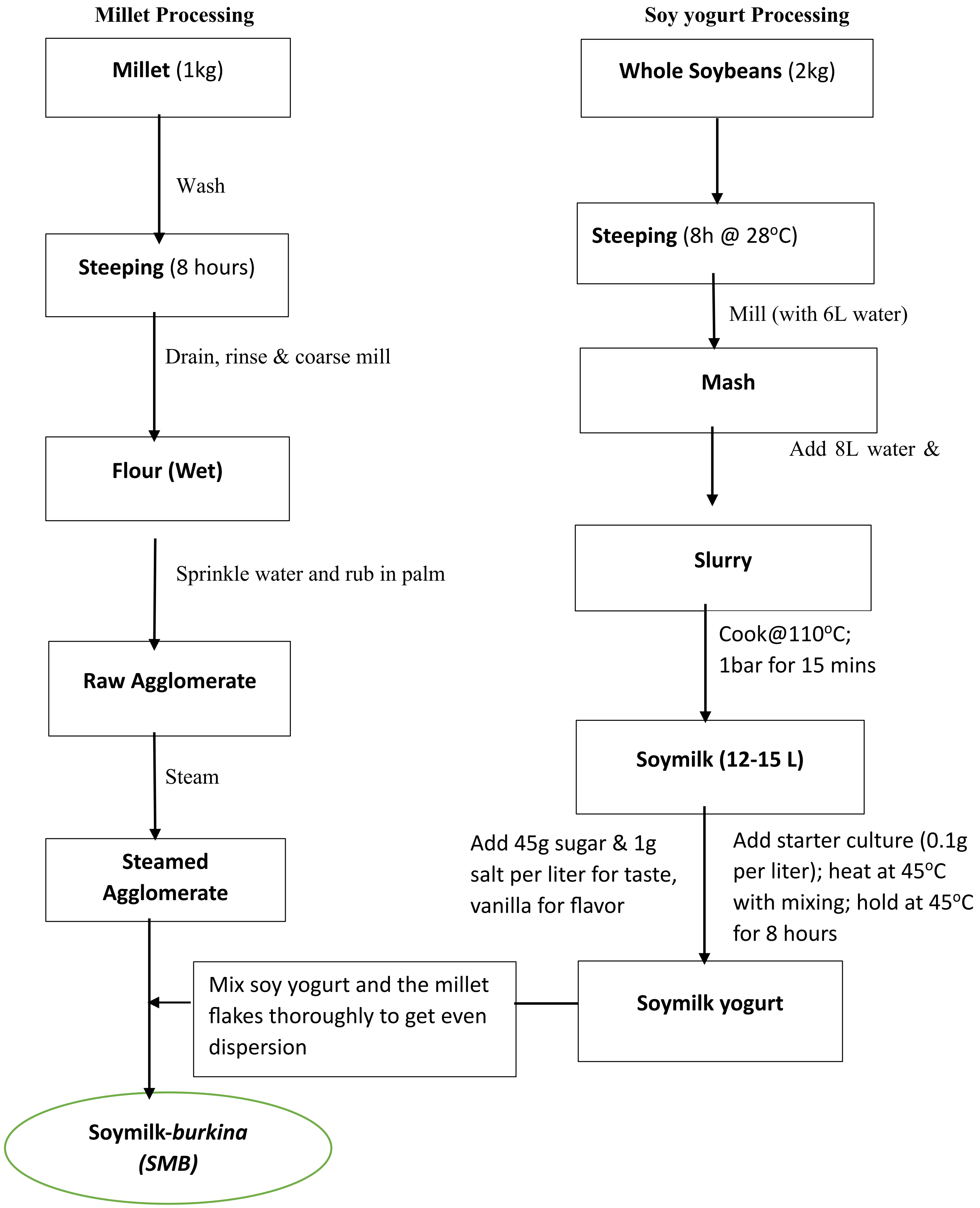


**Supplementary Figure 1. Flow diagram for Soymilk- *burkina* production**

**Supplementary Table 1**: **Table of Illumina sequenced samples used for the analysis with their accession numbers.**

| Accession | Sample Name | SPUID | Organism | Tax ID |
|---|---|---|---|---|
| SAMN35324167 | FLM1-0_S48 | FLM1-0_S48 | human gut metagenome | 408170 |
| SAMN35324168 | FLM1-12_S234 | FLM1-12_S234 | human gut metagenome | 408170 |
| SAMN35324169 | FLM1-3_S11 | FLM1-3_S11 | human gut metagenome | 408170 |
| SAMN35324170 | FLM1-4_S81 | FLM1-4_S81 | human gut metagenome | 408170 |
| SAMN35324171 | FLM1-6_S117 | FLM1-6_S117 | human gut metagenome | 408170 |
| SAMN35324172 | FLM1-8_S157 | FLM1-8_S157 | human gut metagenome | 408170 |
| SAMN35324173 | FLM2-10_S197 | FLM2-10_S197 | human gut metagenome | 408170 |
| SAMN35324174 | FLM2-12_S235 | FLM2-12_S235 | human gut metagenome | 408170 |
| SAMN35324175 | FLM2-3_S12 | FLM2-3_S12 | human gut metagenome | 408170 |
| SAMN35324176 | FLM2-4_S82 | FLM2-4_S82 | human gut metagenome | 408170 |
| SAMN35324177 | FLM2-6_S118 | FLM2-6_S118 | human gut metagenome | 408170 |
| SAMN35324178 | FLM3-0_S49 | FLM3-0_S49 | human gut metagenome | 408170 |
| SAMN35324179 | FLM3-10_S198 | FLM3-10_S198 | human gut metagenome | 408170 |
| SAMN35324180 | FLM3-12_S236 | FLM3-12_S236 | human gut metagenome | 408170 |
| SAMN35324181 | FLM3-3_S13 | FLM3-3_S13 | human gut metagenome | 408170 |
| SAMN35324182 | FLM3-6_S120 | FLM3-6_S120 | human gut metagenome | 408170 |
| SAMN35324183 | FLM3-8_S159 | FLM3-8_S159 | human gut metagenome | 408170 |
| SAMN35324184 | FLM4-0_S50 | FLM4-0_S50 | human gut metagenome | 408170 |
| SAMN35324185 | FLM4-12_S237 | FLM4-12_S237 | human gut metagenome | 408170 |
| SAMN35324186 | FLM4-3_S14 | FLM4-3_S14 | human gut metagenome | 408170 |
| SAMN35324187 | FLM4-6_S121 | FLM4-6_S121 | human gut metagenome | 408170 |
| SAMN35324188 | FLM4-8_S160 | FLM4-8_S160 | human gut metagenome | 408170 |
| SAMN35324189 | FLM5-0_S51 | FLM5-0_S51 | human gut metagenome | 408170 |
| SAMN35324190 | FLM5-12_S238 | FLM5-12_S238 | human gut metagenome | 408170 |
| SAMN35324191 | FLM5-3_S15 | FLM5-3_S15 | human gut metagenome | 408170 |
| SAMN35324192 | FLM5-4_S84 | FLM5-4_S84 | human gut metagenome | 408170 |
| SAMN35324193 | FLM5-6_S122 | FLM5-6_S122 | human gut metagenome | 408170 |
| SAMN35324194 | FLM5-8_S161 | FLM5-8_S161 | human gut metagenome | 408170 |

| | | | | |
|---|---|---|---|---|
| SAMN35324195 | FNP1-0_S41 | FNP1-0_S41 | human gut metagenome | 408170 |
| SAMN35324196 | FNP1-10_S186 | FNP1-10_S186 | human gut metagenome | 408170 |
| SAMN35324197 | FNP1-12_S224 | FNP1-12_S224 | human gut metagenome | 408170 |
| SAMN35324198 | FNP1-3_S1 | FNP1-3_S1 | human gut metagenome | 408170 |
| SAMN35324199 | FNP1-4_S72 | FNP1-4_S72 | human gut metagenome | 408170 |
| SAMN35324200 | FNP1-6_S107 | FNP1-6_S107 | human gut metagenome | 408170 |
| SAMN35324201 | FNP10-0_S47 | FNP10-0_S47 | human gut metagenome | 408170 |
| SAMN35324202 | FNP10-10_S195 | FNP10-10_S195 | human gut metagenome | 408170 |
| SAMN35324203 | FNP10-12_S233 | FNP10-12_S233 | human gut metagenome | 408170 |
| SAMN35324204 | FNP10-3_S10 | FNP10-3_S10 | human gut metagenome | 408170 |
| SAMN35324205 | FNP10-4_S80 | FNP10-4_S80 | human gut metagenome | 408170 |
| SAMN35324206 | FNP10-6_S116 | FNP10-6_S116 | human gut metagenome | 408170 |
| SAMN35324207 | FNP10-8_S156 | FNP10-8_S156 | human gut metagenome | 408170 |
| SAMN35324208 | FNP2-10_S187 | FNP2-10_S187 | human gut metagenome | 408170 |
| SAMN35324209 | FNP2-12_S225 | FNP2-12_S225 | human gut metagenome | 408170 |
| SAMN35324210 | FNP2-3_S2 | FNP2-3_S2 | human gut metagenome | 408170 |
| SAMN35324211 | FNP2-4_S73 | FNP2-4_S73 | human gut metagenome | 408170 |
| SAMN35324212 | FNP2-6_S108 | FNP2-6_S108 | human gut metagenome | 408170 |
| SAMN35324213 | FNP2-8_S148 | FNP2-8_S148 | human gut metagenome | 408170 |
| SAMN35324214 | FNP3-10_S188 | FNP3-10_S188 | human gut metagenome | 408170 |
| SAMN35324215 | FNP3-12_S226 | FNP3-12_S226 | human gut metagenome | 408170 |
| SAMN35324216 | FNP3-3_S3 | FNP3-3_S3 | human gut metagenome | 408170 |
| SAMN35324217 | FNP3-4_S74 | FNP3-4_S74 | human gut metagenome | 408170 |
| SAMN35324218 | FNP3-6_S109 | FNP3-6_S109 | human gut metagenome | 408170 |
| SAMN35324219 | FNP3-8_S149 | FNP3-8_S149 | human gut metagenome | 408170 |
| SAMN35324220 | FNP4-0_S42 | FNP4-0_S42 | human gut metagenome | 408170 |
| SAMN35324221 | FNP4-10_S189 | FNP4-10_S189 | human gut metagenome | 408170 |
| SAMN35324222 | FNP4-12_S227 | FNP4-12_S227 | human gut metagenome | 408170 |
| SAMN35324223 | FNP4-3_S4 | FNP4-3_S4 | human gut metagenome | 408170 |
| SAMN35324224 | FNP4-4_S75 | FNP4-4_S75 | human gut metagenome | 408170 |
| SAMN35324225 | FNP4-6_S110 | FNP4-6_S110 | human gut metagenome | 408170 |
| SAMN35324226 | FNP4-8_S150 | FNP4-8_S150 | human gut metagenome | 408170 |

| | | | | |
|---|---|---|---|---|
| SAMN35324227 | FNP5-0_S43 | FNP5-0_S43 | human gut metagenome | 408170 |
| SAMN35324228 | FNP5-10_S190 | FNP5-10_S190 | human gut metagenome | 408170 |
| SAMN35324229 | FNP5-12_S228 | FNP5-12_S228 | human gut metagenome | 408170 |
| SAMN35324230 | FNP5-3_S5 | FNP5-3_S5 | human gut metagenome | 408170 |
| SAMN35324231 | FNP5-4_S76 | FNP5-4_S76 | human gut metagenome | 408170 |
| SAMN35324232 | FNP5-6_S111 | FNP5-6_S111 | human gut metagenome | 408170 |
| SAMN35324233 | FNP5-8_S151 | FNP5-8_S151 | human gut metagenome | 408170 |
| SAMN35324234 | FNP6-0_S44 | FNP6-0_S44 | human gut metagenome | 408170 |
| SAMN35324235 | FNP6-10_S191 | FNP6-10_S191 | human gut metagenome | 408170 |
| SAMN35324236 | FNP6-12_S229 | FNP6-12_S229 | human gut metagenome | 408170 |
| SAMN35324237 | FNP6-4_S77 | FNP6-4_S77 | human gut metagenome | 408170 |
| SAMN35324238 | FNP6-6_S112 | FNP6-6_S112 | human gut metagenome | 408170 |
| SAMN35324239 | FNP6-8_S152 | FNP6-8_S152 | human gut metagenome | 408170 |
| SAMN35324240 | FNP7-10_S192 | FNP7-10_S192 | human gut metagenome | 408170 |
| SAMN35324241 | FNP7-12_S230 | FNP7-12_S230 | human gut metagenome | 408170 |
| SAMN35324242 | FNP7-3_S7 | FNP7-3_S7 | human gut metagenome | 408170 |
| SAMN35324243 | FNP7-4_S78 | FNP7-4_S78 | human gut metagenome | 408170 |
| SAMN35324244 | FNP7-6_S113 | FNP7-6_S113 | human gut metagenome | 408170 |
| SAMN35324245 | FNP7-8_S153 | FNP7-8_S153 | human gut metagenome | 408170 |
| SAMN35324246 | FNP8-0_S45 | FNP8-0_S45 | human gut metagenome | 408170 |
| SAMN35324247 | FNP8-10_S193 | FNP8-10_S193 | human gut metagenome | 408170 |
| SAMN35324248 | FNP8-12_S231 | FNP8-12_S231 | human gut metagenome | 408170 |
| SAMN35324249 | FNP8-3_S8 | FNP8-3_S8 | human gut metagenome | 408170 |
| SAMN35324250 | FNP8-6_S114 | FNP8-6_S114 | human gut metagenome | 408170 |
| SAMN35324251 | FNP9-0_S46 | FNP9-0_S46 | human gut metagenome | 408170 |
| SAMN35324252 | FNP9-10_S194 | FNP9-10_S194 | human gut metagenome | 408170 |
| SAMN35324253 | FNP9-12_S232 | FNP9-12_S232 | human gut metagenome | 408170 |
| SAMN35324254 | FNP9-3_S9 | FNP9-3_S9 | human gut metagenome | 408170 |
| SAMN35324255 | FNP9-4_S79 | FNP9-4_S79 | human gut metagenome | 408170 |
| SAMN35324256 | FNP9-6_S115 | FNP9-6_S115 | human gut metagenome | 408170 |
| SAMN35324257 | FPW1-10_S201 | FPW1-10_S201 | human gut metagenome | 408170 |
| SAMN35324258 | FPW1-12_S239 | FPW1-12_S239 | human gut metagenome | 408170 |

| | | | | |
|---|---|---|---|---|
| SAMN35324259 | FPW1-3_S16 | FPW1-3_S16 | human gut metagenome | 408170 |
| SAMN35324260 | FPW1-4_S85 | FPW1-4_S85 | human gut metagenome | 408170 |
| SAMN35324261 | FPW1-8_S162 | FPW1-8_S162 | human gut metagenome | 408170 |
| SAMN35324262 | FPW2-0_S52 | FPW2-0_S52 | human gut metagenome | 408170 |
| SAMN35324263 | FPW2-10_S202 | FPW2-10_S202 | human gut metagenome | 408170 |
| SAMN35324264 | FPW2-12_S240 | FPW2-12_S240 | human gut metagenome | 408170 |
| SAMN35324265 | FPW2-4_S86 | FPW2-4_S86 | human gut metagenome | 408170 |
| SAMN35324266 | FPW2-6_S124 | FPW2-6_S124 | human gut metagenome | 408170 |
| SAMN35324267 | FPW2-8_S163 | FPW2-8_S163 | human gut metagenome | 408170 |
| SAMN35324268 | FPW3-0_S53 | FPW3-0_S53 | human gut metagenome | 408170 |
| SAMN35324269 | FPW3-10_S203 | FPW3-10_S203 | human gut metagenome | 408170 |
| SAMN35324270 | FPW3-12_S241 | FPW3-12_S241 | human gut metagenome | 408170 |
| SAMN35324271 | FPW3-3_S18 | FPW3-3_S18 | human gut metagenome | 408170 |
| SAMN35324272 | FPW3-6_S125 | FPW3-6_S125 | human gut metagenome | 408170 |
| SAMN35324273 | FPW3-8_S164 | FPW3-8_S164 | human gut metagenome | 408170 |
| SAMN35324274 | FPW4-0_S54 | FPW4-0_S54 | human gut metagenome | 408170 |
| SAMN35324275 | FPW4-10_S204 | FPW4-10_S204 | human gut metagenome | 408170 |
| SAMN35324276 | FPW4-12_S242 | FPW4-12_S242 | human gut metagenome | 408170 |
| SAMN35324277 | FPW4-3_S19 | FPW4-3_S19 | human gut metagenome | 408170 |
| SAMN35324278 | FPW4-4_S87 | FPW4-4_S87 | human gut metagenome | 408170 |
| SAMN35324279 | FPW4-6_S126 | FPW4-6_S126 | human gut metagenome | 408170 |
| SAMN35324280 | FPW4-8_S165 | FPW4-8_S165 | human gut metagenome | 408170 |
| SAMN35324281 | FPW5-0_S55 | FPW5-0_S55 | human gut metagenome | 408170 |
| SAMN35324282 | FPW5-10_S205 | FPW5-10_S205 | human gut metagenome | 408170 |
| SAMN35324283 | FPW5-12_S243 | FPW5-12_S243 | human gut metagenome | 408170 |
| SAMN35324284 | FPW5-3_S20 | FPW5-3_S20 | human gut metagenome | 408170 |
| SAMN35324285 | FPW5-4_S88 | FPW5-4_S88 | human gut metagenome | 408170 |
| SAMN35324286 | FPW5-6_S127 | FPW5-6_S127 | human gut metagenome | 408170 |
| SAMN35324287 | FPW5-8_S166 | FPW5-8_S166 | human gut metagenome | 408170 |
| SAMN35324288 | UFLM1-0_S63 | UFLM1-0_S63 | human gut metagenome | 408170 |
| SAMN35324289 | UFLM1-12_S253 | UFLM1-12_S253 | human gut metagenome | 408170 |
| SAMN35324290 | UFLM1-3_S31 | UFLM1-3_S31 | human gut metagenome | 408170 |

| | | | | |
|---|---|---|---|---|
| SAMN35324291 | UFLM1-4_S98 | UFLM1-4_S98 | human gut metagenome | 408170 |
| SAMN35324292 | UFLM1-6_S137 | UFLM1-6_S137 | human gut metagenome | 408170 |
| SAMN35324293 | UFLM1-8_S176 | UFLM1-8_S176 | human gut metagenome | 408170 |
| SAMN35324294 | UFLM2-0_S64 | UFLM2-0_S64 | human gut metagenome | 408170 |
| SAMN35324295 | UFLM2-10_S214 | UFLM2-10_S214 | human gut metagenome | 408170 |
| SAMN35324296 | UFLM2-3_S32 | UFLM2-3_S32 | human gut metagenome | 408170 |
| SAMN35324297 | UFLM2-4_S99 | UFLM2-4_S99 | human gut metagenome | 408170 |
| SAMN35324298 | UFLM3-10_S215 | UFLM3-10_S215 | human gut metagenome | 408170 |
| SAMN35324299 | UFLM3-12_S254 | UFLM3-12_S254 | human gut metagenome | 408170 |
| SAMN35324300 | UFLM3-3_S33 | UFLM3-3_S33 | human gut metagenome | 408170 |
| SAMN35324301 | UFLM3-4_S100 | UFLM3-4_S100 | human gut metagenome | 408170 |
| SAMN35324302 | UFLM3-6_S138 | UFLM3-6_S138 | human gut metagenome | 408170 |
| SAMN35324303 | UFLM4-0_S65 | UFLM4-0_S65 | human gut metagenome | 408170 |
| SAMN35324304 | UFLM4-10_S216 | UFLM4-10_S216 | human gut metagenome | 408170 |
| SAMN35324305 | UFLM4-12_S255 | UFLM4-12_S255 | human gut metagenome | 408170 |
| SAMN35324306 | UFLM4-3_S34 | UFLM4-3_S34 | human gut metagenome | 408170 |
| SAMN35324307 | UFLM4-6_S139 | UFLM4-6_S139 | human gut metagenome | 408170 |
| SAMN35324308 | UFLM4-8_S178 | UFLM4-8_S178 | human gut metagenome | 408170 |
| SAMN35324309 | UFLM5-0_S66 | UFLM5-0_S66 | human gut metagenome | 408170 |
| SAMN35324310 | UFLM5-10_S217 | UFLM5-10_S217 | human gut metagenome | 408170 |
| SAMN35324311 | UFLM5-12_S256 | UFLM5-12_S256 | human gut metagenome | 408170 |
| SAMN35324312 | UFLM5-3_S35 | UFLM5-3_S35 | human gut metagenome | 408170 |
| SAMN35324313 | UFLM5-4_S102 | UFLM5-4_S102 | human gut metagenome | 408170 |
| SAMN35324314 | UFLM5-6_S140 | UFLM5-6_S140 | human gut metagenome | 408170 |
| SAMN35324315 | UFLM5-8_S179 | UFLM5-8_S179 | human gut metagenome | 408170 |
| SAMN35324316 | UFLM6-0_S67 | UFLM6-0_S67 | human gut metagenome | 408170 |
| SAMN35324317 | UFLM6-10_S218 | UFLM6-10_S218 | human gut metagenome | 408170 |
| SAMN35324318 | UFLM6-12_S257 | UFLM6-12_S257 | human gut metagenome | 408170 |
| SAMN35324319 | UFLM6-4_S103 | UFLM6-4_S103 | human gut metagenome | 408170 |
| SAMN35324320 | UFLM6-6_S141 | UFLM6-6_S141 | human gut metagenome | 408170 |
| SAMN35324321 | UFLM6-8_S180 | UFLM6-8_S180 | human gut metagenome | 408170 |
| SAMN35324322 | UFLM7-0_S68 | UFLM7-0_S68 | human gut metagenome | 408170 |

| | | | | |
|---|---|---|---|---|
| SAMN35324323 | UFLM7-10_S219 | UFLM7-10_S219 | human gut metagenome | 408170 |
| SAMN35324324 | UFLM7-12_S258 | UFLM7-12_S258 | human gut metagenome | 408170 |
| SAMN35324325 | UFLM7-4_S104 | UFLM7-4_S104 | human gut metagenome | 408170 |
| SAMN35324326 | UFLM7-6_S142 | UFLM7-6_S142 | human gut metagenome | 408170 |
| SAMN35324327 | UFLM7-8_S181 | UFLM7-8_S181 | human gut metagenome | 408170 |
| SAMN35324328 | UFNP1-0_S56 | UFNP1-0_S56 | human gut metagenome | 408170 |
| SAMN35324329 | UFNP1-10_S206 | UFNP1-10_S206 | human gut metagenome | 408170 |
| SAMN35324330 | UFNP1-12_S244 | UFNP1-12_S244 | human gut metagenome | 408170 |
| SAMN35324331 | UFNP1-3_S21 | UFNP1-3_S21 | human gut metagenome | 408170 |
| SAMN35324332 | UFNP1-4_S89 | UFNP1-4_S89 | human gut metagenome | 408170 |
| SAMN35324333 | UFNP1-6_S128 | UFNP1-6_S128 | human gut metagenome | 408170 |
| SAMN35324334 | UFNP1-8_S167 | UFNP1-8_S167 | human gut metagenome | 408170 |
| SAMN35324335 | UFNP10-0_S62 | UFNP10-0_S62 | human gut metagenome | 408170 |
| SAMN35324336 | UFNP10-10_S213 | UFNP10-10_S213 | human gut metagenome | 408170 |
| SAMN35324337 | UFNP10-12_S252 | UFNP10-12_S252 | human gut metagenome | 408170 |
| SAMN35324338 | UFNP10-3_S30 | UFNP10-3_S30 | human gut metagenome | 408170 |
| SAMN35324339 | UFNP10-4_S97 | UFNP10-4_S97 | human gut metagenome | 408170 |
| SAMN35324340 | UFNP10-6_S136 | UFNP10-6_S136 | human gut metagenome | 408170 |
| SAMN35324341 | UFNP10-8_S175 | UFNP10-8_S175 | human gut metagenome | 408170 |
| SAMN35324342 | UFNP2-12_S245 | UFNP2-12_S245 | human gut metagenome | 408170 |
| SAMN35324343 | UFNP2-4_S90 | UFNP2-4_S90 | human gut metagenome | 408170 |
| SAMN35324344 | UFNP2-6_S129 | UFNP2-6_S129 | human gut metagenome | 408170 |
| SAMN35324345 | UFNP2-8_S168 | UFNP2-8_S168 | human gut metagenome | 408170 |
| SAMN35324346 | UFNP3-0_S57 | UFNP3-0_S57 | human gut metagenome | 408170 |
| SAMN35324347 | UFNP3-10_S207 | UFNP3-10_S207 | human gut metagenome | 408170 |
| SAMN35324348 | UFNP3-12_S246 | UFNP3-12_S246 | human gut metagenome | 408170 |
| SAMN35324349 | UFNP3-3_S23 | UFNP3-3_S23 | human gut metagenome | 408170 |
| SAMN35324350 | UFNP3-4_S91 | UFNP3-4_S91 | human gut metagenome | 408170 |
| SAMN35324351 | UFNP3-6_S130 | UFNP3-6_S130 | human gut metagenome | 408170 |
| SAMN35324352 | UFNP3-8_S169 | UFNP3-8_S169 | human gut metagenome | 408170 |
| SAMN35324353 | UFNP4-0_S58 | UFNP4-0_S58 | human gut metagenome | 408170 |

| | | | | |
|---|---|---|---|---|
| SAMN35324354 | UFNP4-10_S208 | UFNP4-10_S208 | human gut metagenome | 408170 |
| SAMN35324355 | UFNP4-12_S247 | UFNP4-12_S247 | human gut metagenome | 408170 |
| SAMN35324356 | UFNP4-3_S24 | UFNP4-3_S24 | human gut metagenome | 408170 |
| SAMN35324357 | UFNP4-4_S92 | UFNP4-4_S92 | human gut metagenome | 408170 |
| SAMN35324358 | UFNP4-6_S131 | UFNP4-6_S131 | human gut metagenome | 408170 |
| SAMN35324359 | UFNP4-8_S170 | UFNP4-8_S170 | human gut metagenome | 408170 |
| SAMN35324360 | UFNP5-0_S59 | UFNP5-0_S59 | human gut metagenome | 408170 |
| SAMN35324361 | UFNP5-3_S25 | UFNP5-3_S25 | human gut metagenome | 408170 |
| SAMN35324362 | UFNP6-0_S60 | UFNP6-0_S60 | human gut metagenome | 408170 |
| SAMN35324363 | UFNP6-10_S209 | UFNP6-10_S209 | human gut metagenome | 408170 |
| SAMN35324364 | UFNP6-12_S248 | UFNP6-12_S248 | human gut metagenome | 408170 |
| SAMN35324365 | UFNP6-3_S26 | UFNP6-3_S26 | human gut metagenome | 408170 |
| SAMN35324366 | UFNP6-4_S93 | UFNP6-4_S93 | human gut metagenome | 408170 |
| SAMN35324367 | UFNP6-6_S132 | UFNP6-6_S132 | human gut metagenome | 408170 |
| SAMN35324368 | UFNP6-8_S171 | UFNP6-8_S171 | human gut metagenome | 408170 |
| SAMN35324369 | UFNP7-0_S61 | UFNP7-0_S61 | human gut metagenome | 408170 |
| SAMN35324370 | UFNP7-10_S210 | UFNP7-10_S210 | human gut metagenome | 408170 |
| SAMN35324371 | UFNP7-12_S249 | UFNP7-12_S249 | human gut metagenome | 408170 |
| SAMN35324372 | UFNP7-3_S27 | UFNP7-3_S27 | human gut metagenome | 408170 |
| SAMN35324373 | UFNP7-4_S94 | UFNP7-4_S94 | human gut metagenome | 408170 |
| SAMN35324374 | UFNP7-6_S133 | UFNP7-6_S133 | human gut metagenome | 408170 |
| SAMN35324375 | UFNP7-8_S172 | UFNP7-8_S172 | human gut metagenome | 408170 |
| SAMN35324376 | UFNP8-12_S250 | UFNP8-12_S250 | human gut metagenome | 408170 |
| SAMN35324377 | UFNP8-3_S28 | UFNP8-3_S28 | human gut metagenome | 408170 |
| SAMN35324378 | UFNP8-6_S134 | UFNP8-6_S134 | human gut metagenome | 408170 |
| SAMN35324379 | UFNP8-8_S173 | UFNP8-8_S173 | human gut metagenome | 408170 |
| SAMN35324380 | UFNP9-10_S212 | UFNP9-10_S212 | human gut metagenome | 408170 |
| SAMN35324381 | UFNP9-12_S251 | UFNP9-12_S251 | human gut metagenome | 408170 |
| SAMN35324382 | UFNP9-3_S29 | UFNP9-3_S29 | human gut metagenome | 408170 |
| SAMN35324383 | UFNP9-4_S96 | UFNP9-4_S96 | human gut metagenome | 408170 |
| SAMN35324384 | UFNP9-6_S135 | UFNP9-6_S135 | human gut metagenome | 408170 |
| SAMN35324385 | UFNP9-8_S174 | UFNP9-8_S174 | human gut metagenome | 408170 |

| | | | | |
|---|---|---|---|---|
| SAMN35324386 | UFPW1-10_S220 | UFPW1-10_S220 | human gut metagenome | 408170 |
| SAMN35324387 | UFPW1-12_S259 | UFPW1-12_S259 | human gut metagenome | 408170 |
| SAMN35324388 | UFPW1-3_S36 | UFPW1-3_S36 | human gut metagenome | 408170 |
| SAMN35324389 | UFPW1-6_S143 | UFPW1-6_S143 | human gut metagenome | 408170 |
| SAMN35324390 | UFPW1-8_S182 | UFPW1-8_S182 | human gut metagenome | 408170 |
| SAMN35324391 | UFPW2-0_S69 | UFPW2-0_S69 | human gut metagenome | 408170 |
| SAMN35324392 | UFPW2-10_S221 | UFPW2-10_S221 | human gut metagenome | 408170 |
| SAMN35324393 | UFPW2-12_S260 | UFPW2-12_S260 | human gut metagenome | 408170 |
| SAMN35324394 | UFPW2-3_S37 | UFPW2-3_S37 | human gut metagenome | 408170 |
| SAMN35324395 | UFPW2-4_S105 | UFPW2-4_S105 | human gut metagenome | 408170 |
| SAMN35324396 | UFPW2-6_S144 | UFPW2-6_S144 | human gut metagenome | 408170 |
| SAMN35324397 | UFPW2-8_S183 | UFPW2-8_S183 | human gut metagenome | 408170 |
| SAMN35324398 | UFPW3-10_S222 | UFPW3-10_S222 | human gut metagenome | 408170 |
| SAMN35324399 | UFPW3-12_S261 | UFPW3-12_S261 | human gut metagenome | 408170 |
| SAMN35324400 | UFPW3-3_S38 | UFPW3-3_S38 | human gut metagenome | 408170 |
| SAMN35324401 | UFPW3-4_S106 | UFPW3-4_S106 | human gut metagenome | 408170 |
| SAMN35324402 | UFPW3-6_S145 | UFPW3-6_S145 | human gut metagenome | 408170 |
| SAMN35324403 | UFPW3-8_S184 | UFPW3-8_S184 | human gut metagenome | 408170 |
| SAMN35324404 | UFPW4-0_S70 | UFPW4-0_S70 | human gut metagenome | 408170 |
| SAMN35324405 | UFPW4-10_S223 | UFPW4-10_S223 | human gut metagenome | 408170 |
| SAMN35324406 | UFPW4-12_S262 | UFPW4-12_S262 | human gut metagenome | 408170 |
| SAMN35324407 | UFPW4-3_S39 | UFPW4-3_S39 | human gut metagenome | 408170 |
| SAMN35324408 | UFPW4-4_S119 | UFPW4-4_S119 | human gut metagenome | 408170 |
| SAMN35324409 | UFPW4-6_S146 | UFPW4-6_S146 | human gut metagenome | 408170 |
| SAMN35324410 | UFPW4-8_S185 | UFPW4-8_S185 | human gut metagenome | 408170 |
| SAMN35324411 | UFPW5-0_S71 | UFPW5-0_S71 | human gut metagenome | 408170 |
| SAMN35324412 | UFPW5-3_S40 | UFPW5-3_S40 | human gut metagenome | 408170 |

Key: The sequenced reads are submitted to the NCBI database with study accession number SUB13443825. Individual sample accession numbers for the paired end reads are also provided in the supplementary table 1 above.
